# Novel culture method to diagnose fungal infections with biopsy tissues

**DOI:** 10.1101/2020.01.22.916346

**Authors:** Jiankang Zhao, Yulin Zhang, Yanyan Fan, Zhujia Xiong, Yudi Xia, Xiaohui Zou, Jiajing Han, Binbin Li, Chunlei Wang, Xinmeng Liu, Binghuai Lu, Bin Cao

## Abstract

Biopsy tissue is a difficult sample to gain for the identification of potential pathogen in the clinical laboratory. At present, there are no effective culture methods for small-sized biopsy tissue. In this study, we propose a novel tissue biopsy culture method based on Myco/F lytic liquid culture system, namely, tissue was grinded before injected into culture bottles. More types and numbers of fungi are cultivated using this culture system than sputum or BALF culture. A few of mycobacteria were also detected with ground tissues. The method may be a promising alternative to or supplement for the traditional plate culture in clinical practice.

## Introduction

Infectious diseases are one of the leading causes of morbidity and mortality of patient around the world (1). However, for some slow-growing or fastidious organisms, conventional culture methodology remains face great challenges. For infections caused by bacteria and fungi, precise pathogen identification is critical, and antimicrobial susceptibility testing is important as well for infectious disease clinicians.

Tissue from a sterile site is obtained through invasive aspiration or during a surgical procedure. As a difficult to get and important specimen for the diagnosis of infectious diseases, it should be handled with utmost care and for various microbiological studies. However, culture with a blood agar or/and Sabouraud’s medium often produce a negative result.

The important fungal media in clinical practice are Sabouraud’s medium agar for sputum, bronchoalveolar lavage fluid (BALF) and urine. Furthermore, BACTEC Myco/F Lytic culture system could be used to grow fungi and mycobacteria from blood and sterile body fluid (2, 3). For clinical specimens inoculated on culture agar, only fast-growing fungi with high capacity of sporulation can be detected (4), while slow-growing fungi are usually overlooked (5). Moreover, long-term culture can dry the medium, which is not conducive to fungal growth. This maybe the reasons why only a few species of molds and yeasts are isolated in some clinical microbiology laboratories (6). The Myco/F lytic system is an automated, continuous-monitoring system for fungal and mycobacterial culture, which has been used for detection of bacteria (2), fungi (7, 8) and mycobacteria in blood (2) and sterile body fluid specimens (3, 9). Therefore, considering the small size of the tissue often gained in clinical microbiology laboratory, Myco/F lytic system might be used for the culture of bacteria and fungi after reasonable sample preparation. To date, little study has evaluated the performance of Myco/F lytic system for recovery of fungi and mycobacteria from biopsy issues. In the study, we assessed the efficacy of this culture system in fungal and mycobacterial culture of biopsy tissue.

## Methods

### Study design and population

We designed a retrospective study to assess the fungal and mycobacterial culturing method for tissues of biopsy. The data was recovered from China-Japan Friendship Hospital, Beijing, China, between April 2018 and November 2019. This study enrolled 1261 specimens of biopsy tissues, which came from 1137 patients, between 14 and 98 years old, 741 males and 520 females. Among these, 944 were also sent for detecting *Mycobacterium tuberculosis* (TB) using GeneXpert MTB/RIF assay, and 853 were sent for mycobacteria culture by using BACTEC MGIT 960 system. Fungal culture-positive specimens with novel method based on Myco/F lytic system were categorized into two groups, namely group 1, including simultaneously positive for sputum or BALF culture, and group 2, including specimens negative or not sent for sputum or BALF culture.

### Microbiological culture and GeneXpert MTB/RIF assay

Sputum and BALF culture were carried out using culture agar in line with regular clinical microbiological procedures. Tissue culture was operated using a novel culturing pipeline as shown in Figure 1. Briefly, the biopsy was grinded with a grinder, appropriate amount of sterile saline was added and then transferred to a sterile container with lid, and the suspension was injected into BACTEC Myco/F Lytic (Myco) blood culture bottles and then incubated in BD BACTEC™ blood culture incubator until the positive or negative light alarm (incubation duration 42 days). The positive culture medium was subcultured on blood agar. The clones growing on blood agar were identified using mass spectrometry or sequencing. Moreover, the grinding fluid of biopsy issue were also used for GeneXpert MTB/RIF assay and mycobacteria culture in BACTEC MGIT 960 system, in which samples were determined to be negative if there was no detectable growth after 42 days of incubation.

**Figure 1.**
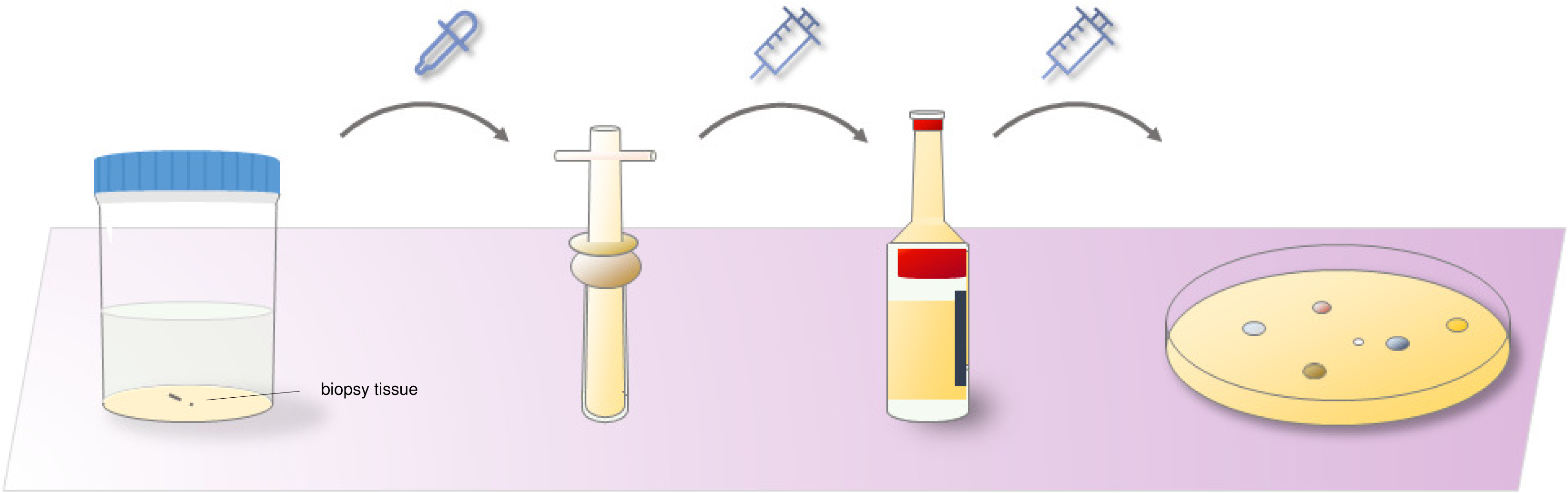
Schematic diagram of fungal or mycobacterial culture of biopsy tissues.

### Definition of infection

To determine whether it is an infection, several indexes were considered in our study, including the pathogenic characteristics of isolated strain, the clinical manifestations of the patient, the sampling method, and the clinical medication. The results were classified into infection, colonization/contamination and unknown.

### Statistical Analysis

Statistical analysis was done by using SPSS Statistics version 21 (IBM). Comparisons of positive rates were done using McNemar’s test. A comparison of normally distributed means was done using an independent samples t test. P value of < 0.05 indicated significant different between groups.

## Results

### Characteristics of samples

Among the 1261 biopsy tissues sent for microbiological assay, 293 specimens (23.24%) were culture positive using novel culture pipeline, including 236 bacteria (18.72%), 39 fungi (3.09%), 16 *Mycobacterium tuberculosis* (1.27%) and 2 non-*Mycobacterium tuberculosis* (0.16%, Figure 2). Furthermore, 25/39 fungi were defined as ID (64.10%), 12/39 were defined as NID (30.77%) and 2/39 were unknown (5.13%).

**Figure 2.**
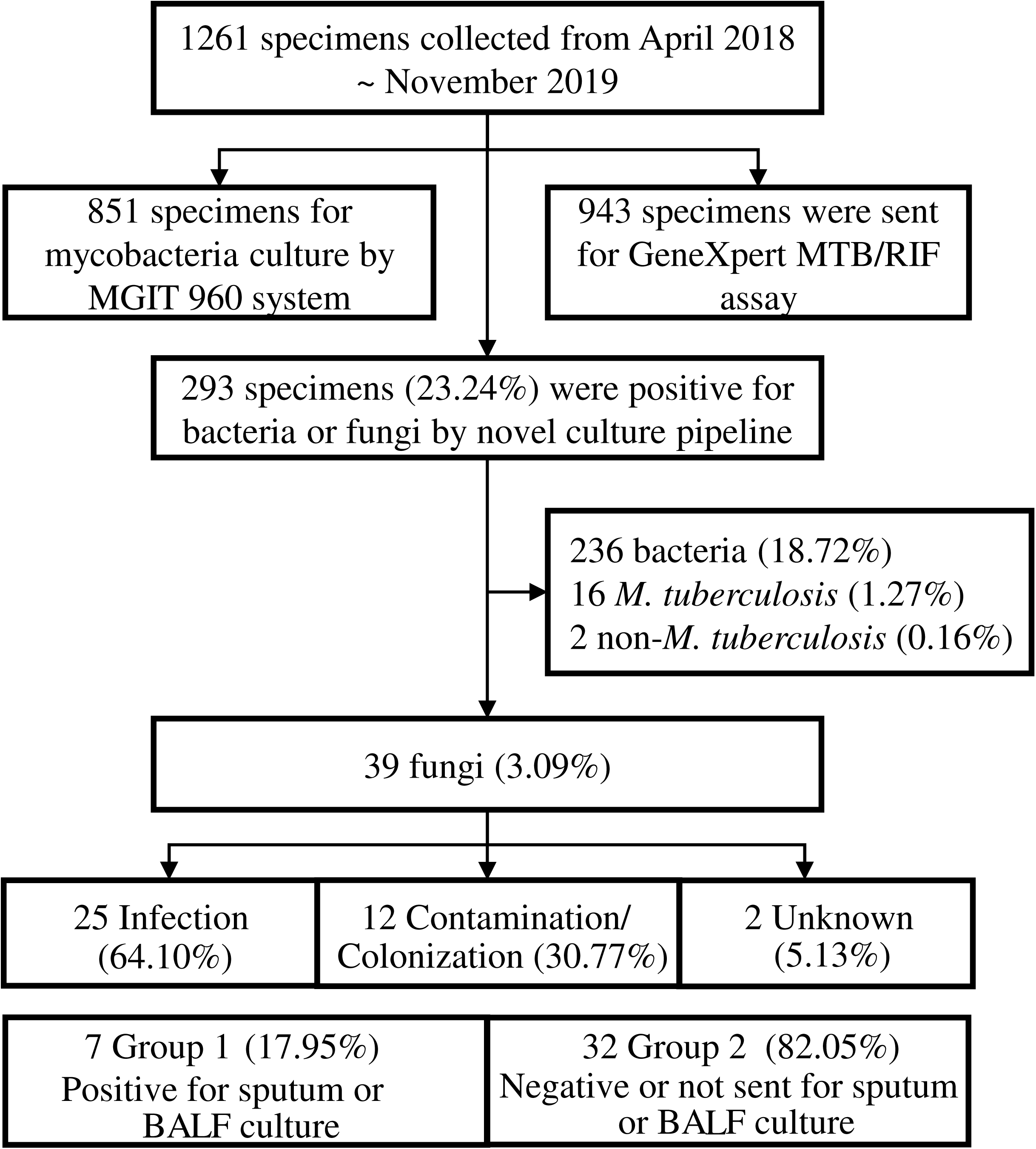
The flow of specimen selection and classification.

### Composition of species in group1 and group2

Among the 39 fungal isolates that were isolated from the biopsy tissues using our method, *Cryptococcus neoformans* (7/39) and *Aspergillus fumigatus* (7/39) were the most isolated species, followed by *Aspergillus flavus* (6/39). *Candida albicans, Aspergillus sydowii, Aspergillus oryzae* and *Schizophyllum commune* all detected in 2 specimens, the other 11 fungal isolates including *Aureobasidium pullulans, Candida glabrata, Candida parapsilosis, Aspergillus niger, Aspergillus versicolor, Trichoderma citrinoviride, Sporothrix globosa, Westerdykella dispersa, Alternaria sp, Penicillium sp* and *Scopulariopsis sp* were detected in one specimen. To assess the effect of novel culturing system in fungal culture, 39 fungal isolates were classified into group 1 and group 2, with 7 (17.95%) and 32 (82.05%) isolates in each group. The 39 isolates consisted of 18 fungal species, isolates in group 1 contained 5 relatively common species including 2 *Aspergillus fumigatus*, 2 *Candida albicans*, 1 *Cryptococcus neoformans*, 1 *Aspergillus niger* and 1 *Aspergillus flavus*. The 32 group 2 isolates belonged to 16 fungal species, and they all had more species in group 2 than group 1 (Figure 3A), although not statistically significant. The infection types for each group was analyzed, specimens in infection, colonization/contamination and unknown of group 1 all had larger amount than that of group 2 (Figure 3B).

**Figure 3.**
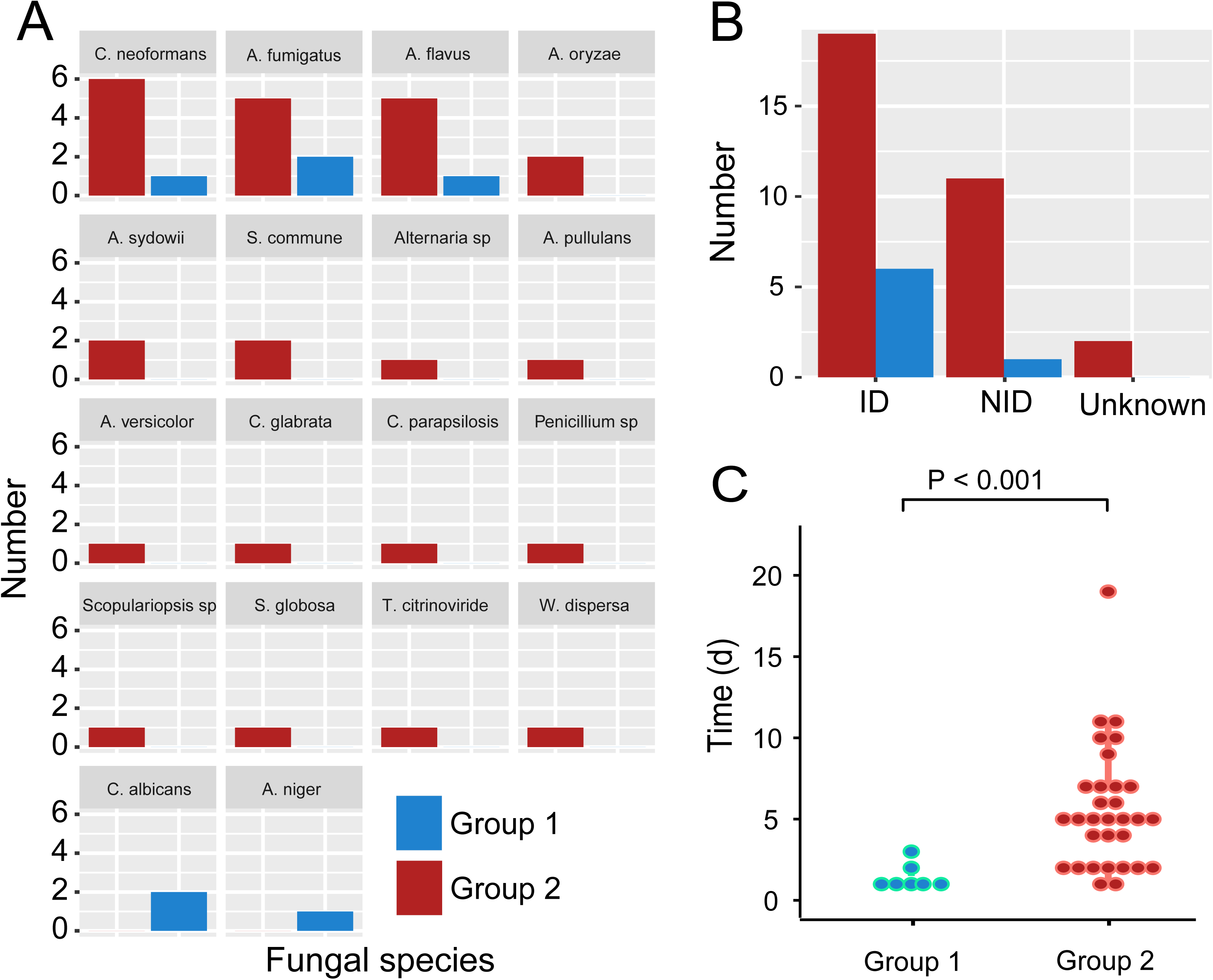
Fungal isolates cultured from biopsy tissues using novel culture pipeline. A. The number of 18 fungal species in group 1 and group 2. B. Number of isolates in group of infection (ID), colonization/contamination (NID) and unknown. C. TTP of 39 fungal isolates in group 1 and group 2. Group 1, including simultaneously positive for sputum or BALF culture, and group 2, including specimens negative or not sent for sputum or BALF culture.

### Time to positivity (TTP)

The TTP of the 39 fungal positive specimens in blood culture incubator was assessed. The median value of TTP was 5 days (1-23 days). 32 specimens in group 2 had a longer incubation time (1-23 days, median value of TTP was 5 days) than 7 specimens in group 1 (1-3 days, median value of TTP 1 days). Statistically significant differences were observed between the two groups with P < 0.001 (Figure 3C).

### Detection of *M. tuberculosis* and non-*M. tuberculosis*

Three methods were used for detection of *M. tuberculosis* in biopsy tissues, namely GeneXpert MTB/RIF assay, BACTEC MGIT 960 system and novel culture pipeline based on BACTEC Myco/F Lytic culture system. Pairwise comparative analysis of the three methods was conducted. A total of 943 specimens were sent for GeneXpert MTB/RIF assay and mycobacterial culture using novel culture system at the same time. A total of 12 *M. tuberculosis* (1.27%) were detected with novel culture system, 54 (5.73%) were *M. tuberculosis* positive for GeneXpert assay, and 7 (0.74%) were *M. tuberculosis* positive for both methods. For detection of *M. tuberculosis*, GeneXpert MTB/RIF assay had a significantly higher positive rate with P value < 0.001 (Table 1). Totally 851 specimens were sent for mycobacterial culture with two different liquid media simultaneously, BACTEC MGIT 960 system and novel culture method. 48 *M. tuberculosis* (5.64%) were culture positive using MGIT 960 system, in which 11 (1.29%) were also detected with method based on Myco/F Lytic culture system (Table 2). Positive rate of BACTEC MGIT 960 system was also significantly higher than novel culture system with P value < 0.001. Furthermore, GeneXpert MTB/RIF assay and BACTEC MGIT 960 system were also compared for detection of *M. tuberculosis*. No significant difference in positive rate was found between the two methods.

**Table 1.** Comparison of *M. tuberculosis* recovery in biopsy tissues for GeneXpert versus novel culture method based on Myco/F lytic culture system.

| GeneXpert | Myco/F lytic system |  |  |
| --- | --- | --- | --- |
|  | Positive | Negative | Total |
| Positive | 7(0.74%) | 47(4.98%) | 54(5.73%) |
| Negative | 5(0.53%) | 884(93.74%) | 889(94.27%) |
| Total | 12(1.27%) | 931(98.73%) | 943(100%) |

**Table 2.** Comparison of *M. tuberculosis* recovery in biopsy tissues for MGIT 960 versus novel culture method based on Myco/F lytic culture system.

| MGIT 960 | Myco/F lytic system |  |  |
| --- | --- | --- | --- |
|  | Positive | Negative | Total |
| Positive | 11(1.29%) | 37(4.35%) | 48(5.64%) |
| Negative | 1(0.12%) | 802(94.24%) | 803(94.36%) |
| Total | 12(1.41%) | 839(98.59%) | 851(100%) |

Non-*M. tuberculosis* is a common pathogen. The BACTEC MGIT 960 system is the main detection method for this pathogen. Benefit from the ability to detect *Mycobacterium* of BACTEC Myco/F Lytic (Myco) blood culture bottles, 2 non-*M. tuberculosis* were detected using novel culture system. Meanwhile, 8 non-*M. tuberculosis* were culture positive with MGIT 960. These two methods had significant difference in positive rate (P =0.031, Table 3).

**Table 3.** Comparison of non-*M. tuberculosis* recovery in biopsy tissues for MGIT 960 versus novel culture method based on Myco/F lytic culture system.

| MGIT 960 | Myco/F lytic system |  |  |
| --- | --- | --- | --- |
|  | Positive | Negative | Total |
| Positive | 2 | 6 | 8 |
| Negative | 0 | 843 | 843 |
| Total | 2 | 849 | 851 |

## Discussion

Microbiological culture, identification and drug sensitivity tests are the standard procedure for clinical pathogen detection, culture is the first and key step for most organisms. Among the 1261 biopsy specimens between April 8, 2018 and November 1, 2019, 293/1261 (23.24%) were positive for bacteria, fungi or mycobacteria. Considering that bacteria can be easily cultured using traditional culture medium, only fungi and mycobacteria were included in the next analysis. Totally 18 different fungal species were cultured, which was much more diverse than previous report (2). *Candida* such as *C. albicans, C. glabrata, C. parapsilosis* and *Aspergillus* such as *A. niger, A. flavus, A. fumigatus* are important clinical strains (10), both of them were isolated using our culturing system. Moreover, 16 *M. tuberculosis* and 2 non-*M. tuberculosis* were also isolated.

Fast and exactly diagnosis of pathogenic microorganisms has great significance for clinical anti-infective therapy. Several factors may infect the incubation time of biopsy tissue in blood culture incubator, such as the initial amount and growth rate of pathogenic microorganisms and the culture condition. The first two factors are usually uncontrollable, but the culture conditions can be optimized. In this novel culture system, Myco/F Lytic (Myco) blood culture bottles provided enough nutrition and time for pathogenic microorganism to grow. A total of 39 fungal isolates were cultured using novel culture pipeline. These isolates were classified into two groups, group 2 (32/39, 82.05%) had much more isolates than group 1 (7/39, 17.95%), which indicated that our novel culture pipeline had advantage for fungal culture than sputum or BALF culture.

For fungal culture of sputum and BALF on culture agar, some slow-growing pathogens cannot be detected. The mean TTP of 7 fungal isolates in group 1 (including 4 *Aspergillus*, 2 *C. albicans* and 1 *C. neoformans*) was 1.43 days, which was similar to previously reported TTP for *C. albicans* (11) and *Aspergillus* (8) in blood. For 32 fungal isolates group 2, the mean TTP was 6.13 days (1-23 days), which seemed too long for fungal culture of sputum and BALF using nutrient agar medium. These isolates consisted of 16 different fungal species. It was our conviction that for fungal culture of biopsy tissues, our novel culture method could be a better choice.

Mycobacterial culture is the gold standard for *M. tuberculosis* diagnosis (12). BACTEC MGIT 960 system is an advanced diagnostic tool commonly used for mycobacteria recovery in clinical laboratories worldwide (13-15). A mixture of antibiotic, including polymyxin B, amphotericin B, nalidixic acid, trimethoprim and azlocillin is added to cultures to decrease contamination rate (16). In addition, account for its high sensitivity and specificity, rapid detection of *M. tuberculosis* and rifampicin resistance, GeneXpert MTB/RIF assay has been widely used in clinical practice (17-20). Myco/F Lytic blood culture bottles was also used for mycobacterial culture. In the present study, a pairwise comparison of the three methods was performed to assess the ability for *M. tuberculosis* detection of biopsy tissues. 943 specimens were sent for GeneXpert assay and mycobacterial culture using novel method simultaneously, GeneXpert (54, 5.73%) had significantly higher positive rate than novel culture method (12, 1.27%) with P value < 0.001. 851 specimens were sent for mycobacterial culture using MGIT 960 and novel culture method base on BACTEC Myco/F Lytic culture system at the meantime, positive rate of MGIT 960 (48. 5.64%) was also higher than novel culture method (12, 1.41%) with P value < 0.001, which was opposite with previous report (3). Another study showed similar positive rate for pleural fluid tuberculosis detected by using MGIT 960 and Myco/F lytic liquid culture system (9). It is generally agreed that the inoculum size of specimen can affect the yield of cultures. In this study, nearly the same volume of specimens was used for culture of mycobacteria. Therefore, specimen volume could not be the determinant for the significant difference between these culture methods. One of the reasons for the significantly lower positive rate of our novel culture pipeline maybe that fast-growing bacteria or fungi mask the growth of mycobacteria, another reason may be the differences in additives between two methods. In any case, above results demonstrated that GeneXpert and MGIT 960 were better methods for detection of *M. tuberculosis*.

Comparison between GeneXpert and MGIT 960 displayed similar positive rate (48/720, 6.67% and 38/720, 5.28%) for detection of *M. tuberculosis*, with no significant difference (P = 0.099). In addition, some specimens were positive for *M. tuberculosis* in only one method of GeneXpert and MGIT 960, this raised the possibility that combination of GeneXpert and MGIT 960 could improve the overall diagnostic yield.

Our study is not without limitations. The major limitation is the small sample size of fungal positive specimens, which is partly because that fungal infection is not as much as bacterial infection. Another reason maybe that some clinicians don’t know the biopsy tissues can be used for fungal culture. In addition, the 39 fungal isolates were grouped into three groups of infection, colonization/contamination and unknown. Isolates in the second group was not low (12, 30.77%), indicating that more standardized specimen processing methods are needed.

In summary, we propose a novel culture process for biopsy tissues, which is very promising for various fungal culture.

